# Sequential Transcriptional Programs of Tissue Residency Drive Human Uterine NK Cell Development

**DOI:** 10.1101/2024.02.29.582684

**Authors:** Morgan Greene, Rebecca Asiimwe, Emma Wright, Markayla Bell, Daniel Epstein, Stefani D. Yates, Brittney Knott, Samantha Fry, Stephanie Clevenger, Jayme Locke, Shawn C. Little, Holly Richter, Aharon G. Freud, Paige M. Porrett

**Author notes:** Equal contributions (Morgan Greene, Rebecca Asiimwe).

## Abstract

Uterine natural killer cells are critical for pregnancy success, but the origin and development of these cells in humans remain unclear. Here we use various single cell approaches to identify the transcriptional programs governing uterine NK cell development in humans. These analyses suggest a developmental continuum which begins with seeding of the endometrium with blood immature CD56^bright^ precursors, evolves through CD56^bright^ endometrial founder NK cells, and ends with tissue resident decidual NK cells during pregnancy which possess non-cytotoxic functions. Our work identifies a role for sequential programs of tissue residency in the differentiation of these cells, as differentiating endometrial tissue resident NK (trNK) cells acquire early and late transcriptional programs of residency which coincide with acquisition of unique non-cytotoxic effector programs. Notably, we identified early residency programs in human endometrial trNKs by expression of *NR4A2*, AP-1 transcription factors, and other immediate early response genes that were shared with CD8 tissue resident memory T cells in mice, suggesting conservation of transcriptional programs of early tissue residency programs across species and cell types. Late residency programs were guided by TGFβ, which promoted expression of various integrins and trNK subset diversification within the non-pregnant endometrium. Altogether, these data identify the molecular foundations for endometrial trNK heterogeneity and suggest that the uterine NK diversity observed during pregnancy is established before embryo implantation and intimately tied to residency programming.

## INTRODUCTION

Natural killer (NK) cells, integral to the innate immune response, constitute 5-15% of circulating lymphocytes^1^ and possess potent cytotoxic and pro-inflammatory functions without the need for prior sensitization^2,3^. These properties not only render them critical in combating viral infections, particularly those caused by herpesviruses such as cytomegalovirus^4^, but also underscore their vital role in immune surveillance. Recent insights into the diverse functions of NK cells, especially their involvement in tissue-resident immunity, have significantly enhanced our understanding of their biological roles.

The concept of tissue resident immunity, traditionally associated with T cells, has been extended to NK cells, thereby broadening our perspective on lymphocyte function within specific tissue contexts. For instance, CD8+ tissue-resident memory T cells (Trm) are characterized by distinct surface phenotypes and functionalities^5–8^, with studies revealing early tissue-adapted precursors and unique molecular signatures of residency^9^. Likewise, NK cells, which hold similarity to CD8+ T cells^10^, have been identified across various tissues and exhibit functional diversity beyond their circulatory counterparts^11^.

Studies of tissue resident NK (trNK) cells in secondary lymphoid organs such as the tonsil have significantly broadened our comprehension of NK cell ontogeny^12–14^. Subsequent studies have delineated NK cell effector functionalities in peripheral tissues, including the lung^15,16^ and liver ^17,18^, unveiling their heterogeneity and expanded capabilities beyond cytotoxic effector functions^19^. While these findings confirm site-specific differentiation in trNK cells, the unique adaptations within understudied and specialized sites such as the endometrium, remains uncertain.

Uterine NK cells play critical roles in the physiology of pregnancy. Notably, endometrial NK (eNK) cells are abundant in the decidualizing non-pregnant uterus, while decidual NK (dNK) cells comprise the predominant immune cell type^20–22^ in the initial stages of pregnancy. Their essential roles in facilitating processes like placentation^23,24^ and spiral artery remodeling^25–27^ underscore their significance, yet the specific origins of uterine NK cells within the endometrial context remain elusive. Conceptual models of uterine NK development based largely on studies in mice propose that these cells may emerge from either NK precursors located within the endometrium, such as hematopoietic stem cells (HSCs)^28,29^, or from mature CD56-expressing cells circulating in the blood, particularly CD56^bright^CD16- or the more abundant CD56^dim^CD16+ peripheral blood NK (pbNK) subsets^30,31^. Insights from uterus transplant studies support the peripheral origin of uterine NK cells, as NK cells in endometrial biopsies after transplantation are recipient-derived^32^. Additional studies identified that CD56^dim^CD16+ cells have the potential to acquire a dNK phenotype under the influence of TGFβ^33–36^, gene expression profiles indicate that tissue resident NK cells are more likely derived from the less differentiated pbNK population, such as CD56^bright^CD16-cells^11,19,37–39^.

Given the rapid turnover of tissue-resident NK cells across menstrual cycles, we favored a parsimonious model of uterine NK differentiation in which endometrial NK cells predominantly derive from CD56-expressing subsets in the blood, circumventing the necessity for extensive transcriptional reprogramming. To test this, we compared the transcriptomes of peripheral blood NK cells and endometrial NK cells to identify the specific blood-derived populations contributing to the endometrial NK cell pool. Single-cell RNA sequencing (scRNA-seq) allowed us to delineate gene expression profiles and putative developmental states on a single-cell basis, which permitted the identification of the CD56^bright^ endometrial founder cell population that precedes mature tissue resident NK subsets. Comparison of endometrial NK transcriptomes with reference decidual NK transcriptomes suggested that diversification of the trNK repertoire was complete in the absence of pregnancy, while signals from the decidual microenvironment promoted expansion of specific trNK subsets. This approach, complemented by Single Cell Regulatory Network Inference and Clustering (SCENIC)^40^ analysis and CITE-seq analysis of endometrial NK proteomes and transcriptomes, offers a comprehensive view of uterine NK cell development, including the regulatory programs dictating these diverse cellular states and functions.

## RESULTS

### Tissue resident NK cells in human endometrium derive from CD56^bright^CD16^−^ peripheral blood NKs

To determine the origin of endometrial NK cells from peripheral blood, we conducted scRNA-seq on CD45^+^ cells from matched peripheral blood and a secretory phase endometrial biopsy from a healthy control volunteer. We FACS-sorted CD45^+^ cells from both samples to ensure capture of all NK cells given the variability of CD56 expression, and the secretory phase of the menstrual cycle was targeted for biopsy as NK cells are most abundant at this time^41^. To facilitate comparisons between NK cells derived from the blood and the endometrium, we aggregated peripheral blood mononuclear cells (PBMCs) and CD45^+^ cells enriched from the endometrium using Harmony^42^.

Consistent with existing literature, we identified diverse populations of mature immune cells, including T cells, B cells, ILC3s, and monocytes^43–46^. Expectedly, we did not detect immature immune precursors in any compartment given our lack of enrichment for these rare cell types. Mature NK cells, characterized by the expression of cytotoxic effector genes and distinguished from CD8+ T cells by expression of *NCAM1* (CD56) and *FCGR3A* (CD16), were notably more prevalent in the endometrium than in the peripheral blood. Most NK cells collected from endometrial biopsies exhibited a gene expression pattern indicative of tissue residency, which included expression of the integrin *ITGA1* and reduced expression of circulation-associated genes (e.g., *KLF2*), in line with prior studies of tissue resident lymphocytes^22,47–49^.

To further refine our transcriptional analysis, NK cells from both peripheral blood and the endometrium were re-clustered (Clusters 6+10), revealing minimal transcriptional overlap between pbNK and eNK cells. As in the analysis of endometrial CD45 cells, re-clustering of NK cells revealed that subsets of NK cells were first segregated by location given differential expression of genes related to tissue residency (*ITGA1*) and cytotoxic effector functions (*FCGR3A*) (Fig. 1A-C). Differential expression analysis identified 121 genes upregulated in eNK cells, encompassing *NCAM1*, tissue retention markers (*ITGA1, CD69*), and lymphocyte chemotaxis-associated genes (*CCL3, CCL4, XCL1*), alongside canonical decidual NK cell markers^48^ (Fig. 1D). Conversely, 189 genes were found to be more highly expressed in pbNK cells, including *FCGR3A* and genes linked to cytotoxic effector mechanisms (*FGFBP2, CX3CR1, GZMH*) ^10,50,51^. This analysis further identified NKT cells in both peripheral blood and endometrium(cluster 6), as well as CD56^bright^CD16^−^ NK cells (cluster 7), and CD56^dim^CD16^+^ conventional NK cells (cluster 2; cNK), based on CD3 expression and reference transcriptomes ^51^ (Fig. 1E). Interestingly, approximately 4% of eNK cells shared sufficient transcriptional similarity with peripheral blood CD56^dim^ and CD56^bright^ cells to cluster with these populations, suggesting recent migration to the endometrium without significant differentiation (Fig. 1A&B).

**Fig 1.**
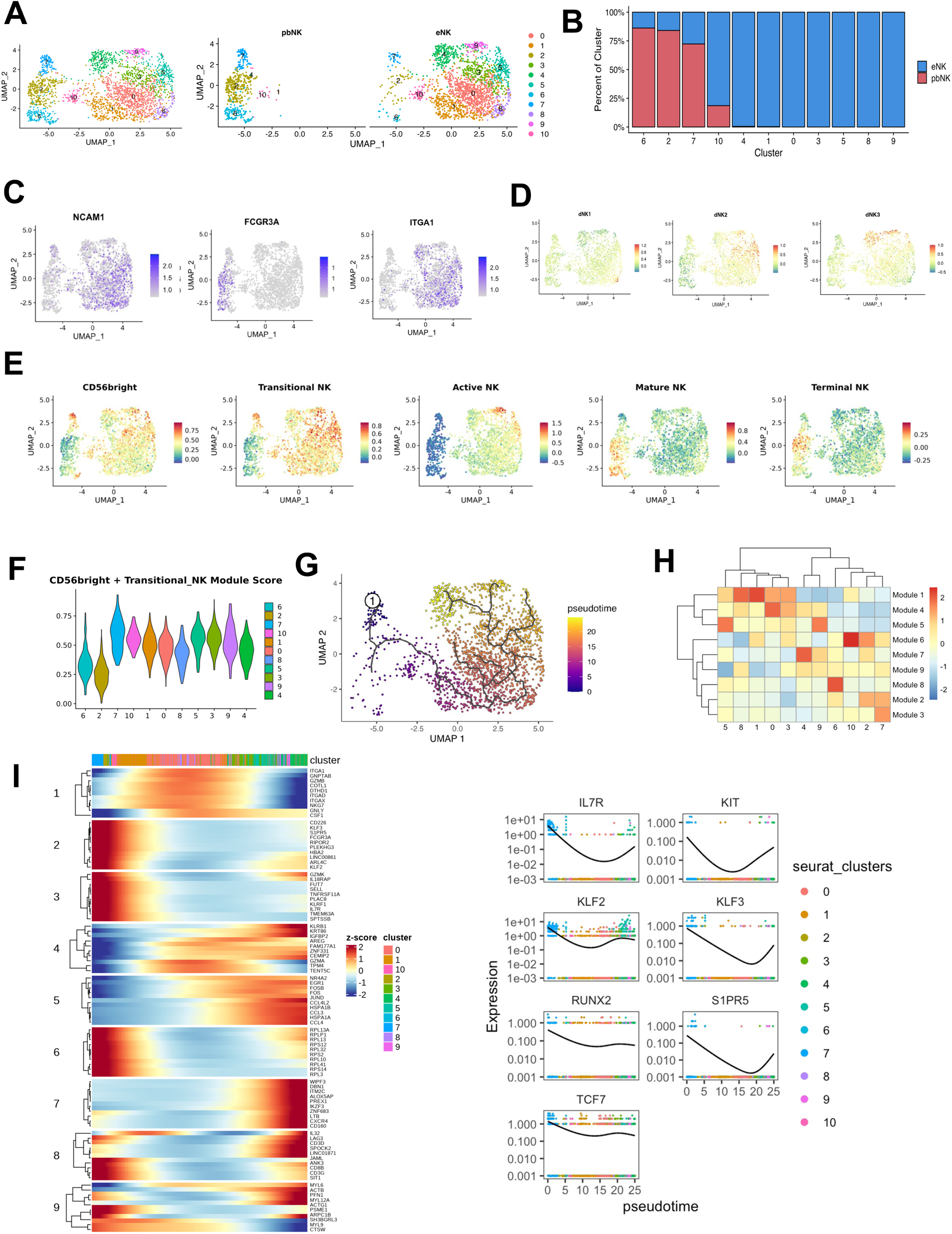
Tissue resident NK cells in human endometrium derive from CD56^bright^CD16-peripheral blood NKs (A) Merged and split UMAP embedding of reclustered NK cells, selected from an aggregation of scRNAseq data from sorted live CD45+ PBMC and endometrial cells from a matched healthy control participant (n=1). (B) Frequency of NK cells derived from peripheral blood (pbNK) or endometrium (eNK) in each cluster shows progressive decrease in pbNK:eNK ratio with clusters 6, 2, and 7 having >50% of pbNK cells. (C) Expression of *NCAM1* and *FCG3RA* identifying circulating NK cell populations (CD56^bright^ CD16^−^, CD56^dim^ CD16^+^) and expression of *ITGA1* identifying tissue resident NK cell populations. Color represents normalized expression. (D-F) Module score of (D) top 15 DEGs expressed by the 3 decidual NK cells subsets, (E) top 50 DEGs of circulating NK cell stages, and (F) combined CD56bright and transitional NK DEGs. (G) Trajectory analysis of eNK development with CD56^bright^ pbNK cells (cluster 7) set as root. (H & I) Heatmap displaying the expression of 9 gene modules identified across trajectory identified in G for each cell cluster; (H) overall module expression (I) expression of module defining genes; color bar above indicates each cells cluster of origin, color scale represents z score

The identification of CD56^bright^ (cluster 7) and CD56^dim^ (cluster 2) cells in the endometrium as potential precursors to mature eNK populations led us to hypothesize that the origins of mature eNK cells could be deduced from shared core transcriptional signatures with putative precursor cells in the blood. Comparative transcriptome analysis between mature eNK cells and reference pbNK CD56^bright^ and CD56^dim^ transcriptomes ^51^ revealed that CD56^bright^ cells from peripheral blood (cluster 7) were the likely precursors to mature eNK populations. This was evidenced by the extensive sharing of the core pbNK CD56^bright^ transcriptional signature with endometrial trNK populations. Nevertheless, there was a reduction of this core pbNK CD56^bright^CD16^−^ signature among mature eNK clusters, suggesting downregulation of this peripheral blood signature as endometrial NK cells differentiated in the tissue from blood CD56^bright^CD16^−^ precursors (i.e., cluster 7 cells derived from the endometrium) (Fig. 1E&F). These data align with the current understanding that peripheral blood CD56^bright^ cells possess stem-like potential and undergo further maturation and terminal differentiation into CD56^dim^ populations^12,13,39,52^.

To further dissect the differentiation process of eNK cells from their progenitors, trajectory analysis was performed with cluster 7 as the root (Fig. 1G). 1,223 genes were differentially expressed along the differentiation trajectory and were organized into 9 modules for in-depth examination (Fig. 1H&I). This analysis identified cluster 10 as a translationally active, intermediate eNK population that bridged the putative CD56^bright^ endometrial founder population (cluster 7) and the rest of the eNK populations, suggesting a phased differentiation pathway (Fig. 1G&H). Notably, the downregulation of genes associated with circulatory potential (*S1PR5, KLF2, and KLF3*) in module 2 and those linked to stemness and immaturity (*TCF7*, *RUNX2, KIT, IL7R*) in module 3 indicates a transition from immature to more differentiated states (Fig. 1I). Collectively, these data suggest that CD56^bright^ pbNK cells seed the endometrium and give rise to distinct tissue resident eNK subsets, while more terminally differentiated blood CD56^dim^ cells, albeit a minority, infiltrate the endometrium and retain their cytotoxic effector functions.

### Initiation of tissue residency by transcriptional reprogramming of founder NK cells

Having identified cluster 7 endometrial NK cells as the candidate tissue-based precursors of mature tissue resident eNK cells, we next aimed to identify the transcriptional programs dictating early tissue residency. As elegant time course studies of lymphocyte tissue residency development in mouse models have identified genes that are upregulated shortly after blood-based precursors enter a tissue^8,9^, we hypothesized that newly arrived cells in the endometrium would exhibit similar gene expression patterns associated with early tissue adaptation. Through differential expression analysis of CD56^bright^ pbNK and eNK cells within cluster 7, we discovered a significant upregulation of immediate early genes in eNK cells, notably within the NR4A family (*NR4A2*) and the AP-1 complex (*FOS, JUN, FOSB*, etc.), relative to their peripheral blood counterparts (Fig. 2A). Importantly, these genes are recognized as essential for establishing tissue residency in murine mucosal CD8 resident memory T cells (Trm) following LCMV infection, with their expression amplifying over time to sustain tissue residency^9,53^. We found that the Trm signature was notably enriched in eNK cells from cluster 7 (Fig 2B), leading us to designate these eNK cells as founder NK cells. Remarkably, this early adaptation signature was not restricted to founder NK cells but was also shared by endometrial NKTs and CD56^dim^CD16^+^ NK cells, suggesting a widespread mechanism of tissue adaptation across lymphocyte populations initiated upon tissue entry (Fig. 2C).

**Fig 2.**
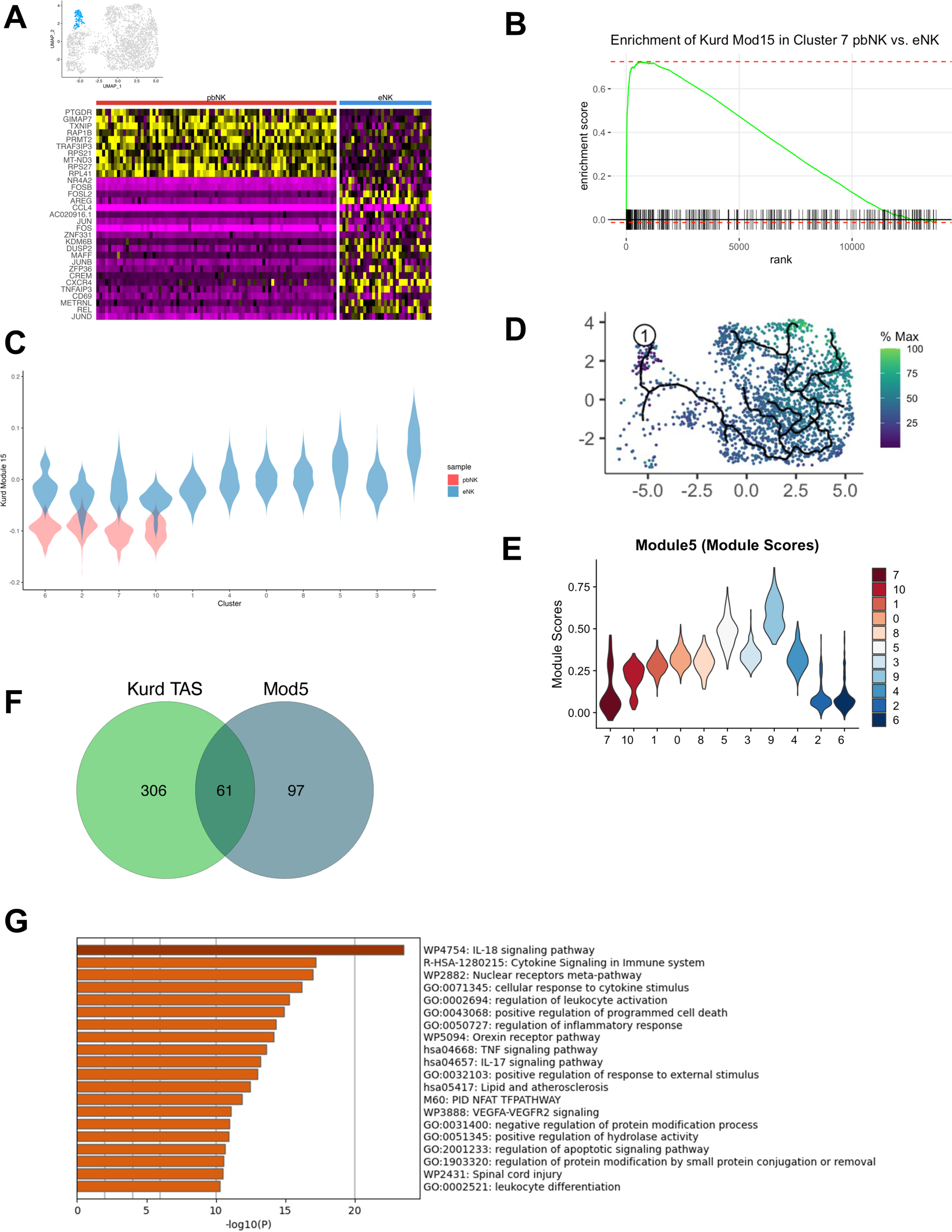
Initiation of tissue residency by transcriptional reprogramming in founder NK cells (A) Heat map of top differentially expressed genes between pbNK and eNK cells within CD56^bright^ cluster (cluster 7) (B) Gene set enrichment analysis (GSEA) showing expression of Trm genesets in eNK (C) Violin plots of relative expression of tissue adaptation module in pbNK (red) vs eNK (blue) (D & E) Visualization of module score 5 identified via trajectory analysis in Fig 1H; (D) expression of module across merged UMAP embedding; color scale represents proportion of genes in the module being expressed (E) Violin plot of module expression across NK clusters (F) Venn diagram showing differentially expressed genes between Kurd module 15 genes and eNK (log2FC > 0.5 and P adj. < 0.05) (G) Pathway analysis of shared genes in (F)

Given the upregulation of early residency genes in murine CD8+ Trm cells over time^9^, we hypothesized that a similar increase in this gene signature would occur in endometrial NK cells as they transitioned into mature tissue resident subsets. Revisiting our trajectory analysis (Fig. 1H), we identified this conserved, early residency gene signature among the genes differentially expressed across the trajectory. Among the 9 gene modules influencing eNK differentiation, we identified the early residency signature in module 5 genes, which consistently increase along the trajectory from cluster 7 to clusters 4 and 9 (Fig. 2D&E). The overlap between trajectory module 5 genes and the early tissue adaptation signature further confirmed this finding, as 51 of the 327 genes in the CD8+ Trm tissue adaptation module were shared with endometrial module 5, including transcription factors required for tissue retention such as *NR4A2, FOS*, and *JUN* (Fig. 2F).. Pathway analysis of trajectory module 5 genes highlighted enrichment for IL-17 and IL-18 cytokine signals, alongside NFAT and VEGF signaling pathways (Fig. 2G), suggesting these signals play a pivotal role in upregulating tissue adaptation genes after endometrial entry. Taken together, these data suggest a model wherein specific cytokine and signaling cues drive early transcriptional shifts foundational for lymphocyte tissue residency. This process, initiated upon the entry of precursor cells into the endometrium, involves a conserved transcriptional response that not only facilitates early tissue adaptation but also steers the differentiation of NK cells into mature tissue resident subsets.

### Tissue residency promotes diversification of endometrial tissue resident NK cells

We next sought to interrogate the transcriptional programs underpinning NK cell heterogeneity. Although decidual NK cell heterogeneity is well appreciated^32,48^ and found in non-pregnant individuals^22,39^, it is unclear when NK cell heterogeneity arises in the endometrium. To this end, we performed scRNA-seq on FACS-sorted CD45+ cells from secretory phase endometrial biopsies of five additional healthy control volunteers and analyzed 14,349 endometrial NK cells that were re-clustered from these aggregated data (n=6 subjects; 41,722 CD45^+^ cells) (Fig. 3A).

**Fig 3.**
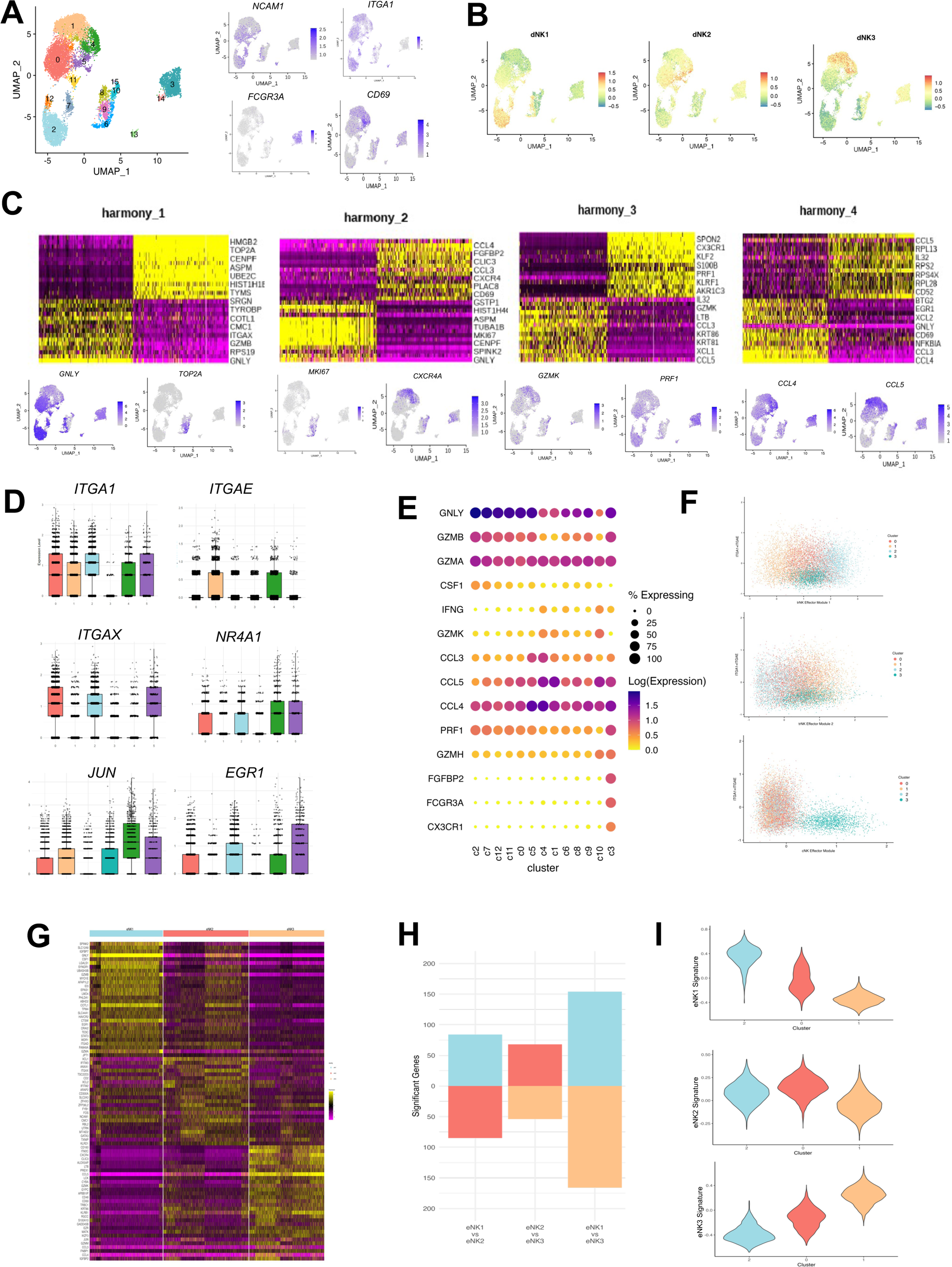
Linked programs of tissue residency and effector function underpin eNK cell diversity. (A) Merged UMAP embedding of 14,349 eNK cells aggregated from 6 healthy-control participants (left) and feature plots highlighting expression of classical NK (*NCAM1, FCGR3A*) and tissue residency (*ITGA1, CD69*) markers. (B) UMAP embedding with cells colored by their average expression of the top 15 differentially expressed genes within the three identified dNK cell subsets. (C) Heatmap presentation of the 15 leading genes from principal component analysis, and feature plots highlighting the expression of the most differentially expressed genes within each PC. (D) Comparison of expression of key genes related to integrin expression (*ITGA1, ITGAE, ITGAX*) and early residency programming (*NR4A1, JUN, EGR1*) across the major NK subsets. (E) Dot plot of specific effector programs within the identified cell clusters. (F) Scatter plot analysis demonstrating the correlation between effector module scores and tissue residency module scores. (G) Heatmap comparison of gene expression profiles across the three principal eNK subsets. (H) Bar graph quantifying the number of differentially expressed genes among the three main NK cell subsets. (I) Violin plots detailing gene expression signatures from the major NK subsets, with overlay on the respective subset profiles.

Our refined analysis identified several subsets of endometrial trNK cells, namely clusters 0 (trNK2), 1 (trNK3), and 2 (trNK1), characterized by the expression of integrin *ITGA1* (CD49a), the absence of *FCGR3A* (CD16), and shared marker genes with decidual NK subsets (Fig. 3A&B)^48^. We resolved additional subsets within the trNK3 and trNK2 clusters which were distinguished by increased expression of AP-1 transcription factors and upregulation of NFkB (cluster 4 & cluster 5, respectively). Multiple subsets of trNK1 cells were also observed (clusters 2, 7, 11, and 12) (Fig. 3A&B), including a cluster not previously described expressing a strong IFN-α gene signature (cluster 11). Distribution of these NK clusters were largely similar across a majority of volunteers, with the notable exception of trNK1, which represented 35% of eNK cells in HC2 compared to the range of 3-12% in the other five subjects.

In dissecting the transcriptional drivers of NK cell diversification, we correlated gene expression patterns to the top harmonized principal components of our integrated dataset. Genes governing cellular proliferation (*MKI67, TOP2A*), effector functions (*PRF1, CCL5, FCGR3A*), and tissue residency or circulation (*CD69, CXCR4, ITGA1, KLF2*) were among the most variable genes in endometrial NK subsets (Fig. 3C). Notably, genes implicated in the early residency programming of founder NK cells were found to contribute to the diversification of endometrial trNK cells, as indicated by their substantial representation in influencing the fourth Harmony-corrected principal component. The distinction between cNK and trNK subsets, as well as among trNK subsets themselves, was underscored by the differential expression of genes related to early residency programming and integrin expression (Fig. 3D).

Further analysis of the transcriptional landscapes revealed distinct effector programs within eNK subsets; cNKs primarily expressed genes related to cytotoxicity, whereas trNK subsets were characterized by the expression of chemokine ligands and cytokine genes, suggesting differentiated roles in immune responses (Fig. 3E). The association between effector functions and tissue residency was further illustrated by correlating integrin gene expression with specific effector gene modules, demonstrating the interconnectedness of these aspects in trNK cells (Fig. 3F). Differential expression analysis among major trNK populations underscored that both effector and residency-defining genes served as key discriminators among these groups (Fig. 3G), with trNK1 and trNK3 showing the greatest divergence, and trNK2 displaying overlapping transcriptomes with both (Fig. 3H&I). This variation supports a model where tissue residency programming plays a pivotal role not just in the differentiation of endometrial trNK cells from their blood-derived precursors but also in the subsequent diversification of trNK subset effector functions within the non-pregnant human endometrium.

### Differential TGFβ imprinting contributes to endometrial trNK diversity

The complexity of integrin expression and the effector function of endometrial trNK cell subsets underscore the sophisticated mechanisms governing their differentiation within the human endometrium. To begin understanding this relationship, we focused on the differential integrin expression identified in our transcriptomic data and prior work. Integrins are crucial for the tissue residency programs of lymphocytes, facilitating tissue retention by mediating interactions with parenchymal cell types^8,19,49,54^. Notably, CD103, expressed on tissue-resident lymphocytes within epithelial tissues of the lung, gut, and skin, serves as the alpha subunit of the αEβ7 heterodimer, interacting with E-cadherin expressed on epithelial cells^8,35,54,55^. Central to this process is the role of TGFβ, a cytokine instrumental in modulating lymphocyte integrin expression, thereby influencing the tissue residency and heterogeneity of lymphocyte populations. Given TGFβ’s critical role and the observed variation in integrin gene expression among endometrial trNK subsets (Fig. 3), we proposed that TGFβ signaling modulation contributes to the diversity of trNK cells.

Building on this premise, we employed CITE-seq analysis on CD45+ cells from three secretory phase endometrial biopsies^56^ to establish a direct connection between the transcriptional programs that govern integrin gene expression and their manifestation as surface proteins. By integrating both proteomic and transcriptomic data with a weighted-nearest neighbor UMAP, we enhanced the resolution of endometrial NK cell clustering beyond what is achievable with RNA-only or antibody-derived tag (ADT; i.e., surface protein) algorithms alone, analyzing a total of 3,957 cells (Fig. 4A)^57^. Consistent with our previous findings, eNK cells formed two major clusters of CD56^bright^CD49a^+^CD16^−^ trNK cells and CD56^dim^CD49a^−^CD16^+^ cNK cells, with a small number of CD56^bright^CD49a^−^CD16^int^ founder cells in proximity to cNK cells. Within the trNK cells, the two predominant clusters exhibited dNK2-like and dNK3-like signatures (Cluster 0 [trNK2] and Cluster 1 [trNK3], respectively), with a small cluster near the trNK2 cells showing a dNK1 transcriptional signature (Cluster 8 [trNK1]) (Fig. 4B).

**Fig 4.**
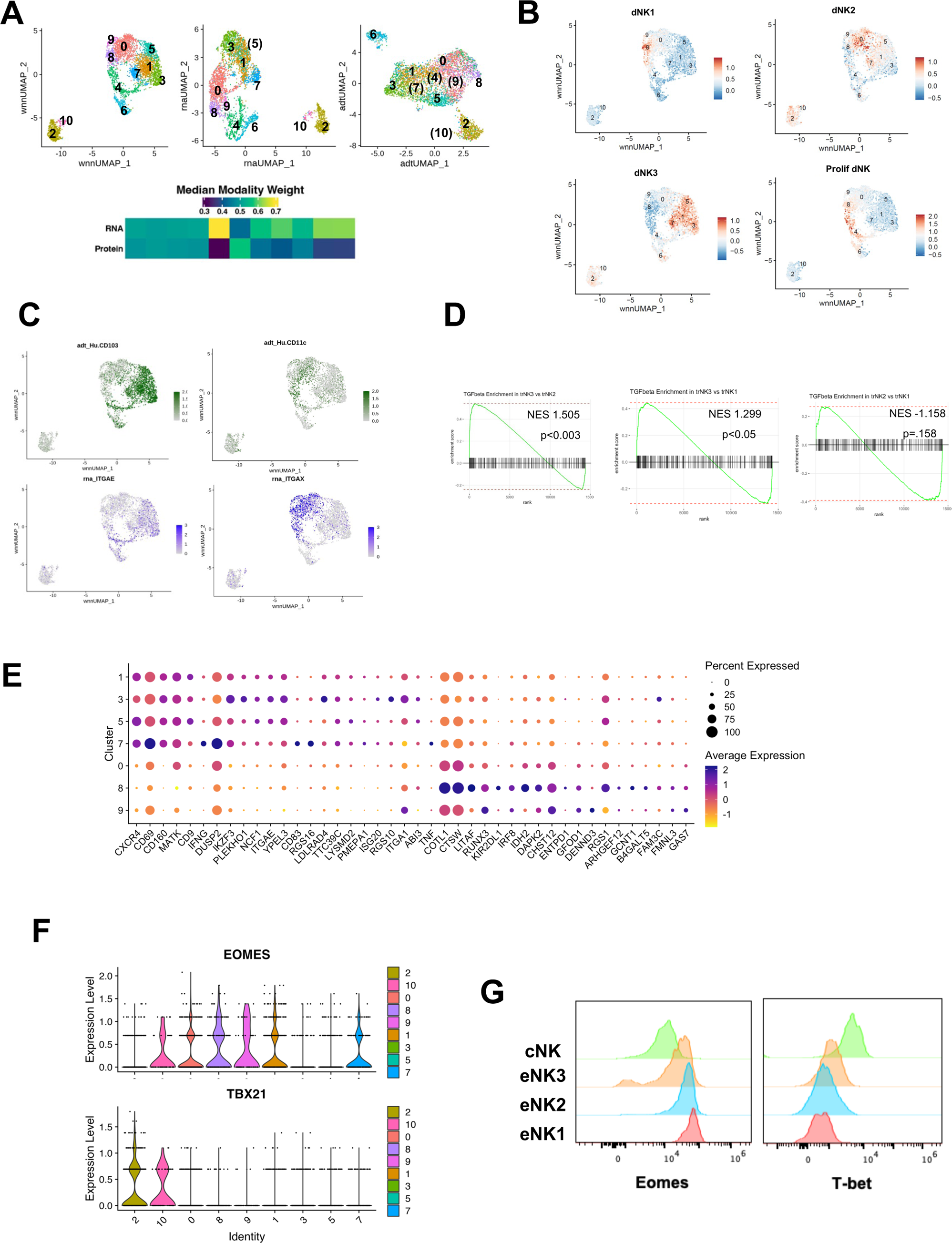
Differential TGFβ imprinting contributes to endometrial trNK diversity. (A) CITE-seq analysis on 3,957 eNK cells from secretory phase endometrial biopsies integrating proteomic and transcriptomic data via weighted-nearest neighbor UMAP. (B) Module score of the 15 most differentially expressed genes within the three identified dNK cell subsets and the proliferating dNK subset. (C) Differential expression of integrins, particularly CD11c (*ITGAX*) and CD103 (*ITGAE*), across trNK cell subsets. (D) Manual curation of genes associated with TGFβ imprinting and GSEA analysis on the major NK subsets. (E) Dot plot of key genes affected by TGFβ imprinting (F) Violin plot comparison of *EOMES (*Eomes) and *TBX21* (T-bet) expression across eNK subsets. (G) Flow cytometry confirmation of the transcription factor expression patterns described in (F).

A key factor in differentiating the principal trNK cell clusters was the expression of integrins, notably CD11c (*ITGAX*) and CD103 (*ITGAE*), which emerged as among the most differentially expressed proteins and genes (Figs. 4C), underscoring the nuanced roles of these molecules in trNK cell identity and function. The differential expression of *ITGB7* on trNK3 cells illustrates that these cells co-express both αEβ7 heterodimer subunits. Although CD103 and ITGB7 were linked to specific transcriptional programs, other integrins like CD49a displayed broader sharing across subsets or varied expression (e.g., CD11c). Despite CD11c’s reciprocal expression on trNK2 and trNK3 cells (Fig. 4C), a subset of CD103+ trNK3 cells also co-expressed CD11c, and CD11c expression did not clearly distinguish between trNK1 and trNK2 cells, highlighting a weak correlation between ITGAX gene expression and CD11c protein presence on trNK1 cells (Figs. 4C). Remarkably, trNK1 cells, displaying the most restricted integrin repertoire among the endometrial trNK subsets, were more accurately differentiated from trNK2 cells through the expression of non-integrin markers such as CD39 and KIR, aligning with previous observations^48^.

In light of the spectrum of integrin expression on trNK subsets, we hypothesized that these subsets were differentially susceptible to TGFβ imprinting. Manual curation of genes associated with TGFβ imprinting and subsequent analysis revealed significant enrichment of TGFβ-responsive genes in trNK3 cells, indicating a more pronounced TGFβ imprinting within this subset (Fig. 4D&E), which was confirmed at the protein level for CD101 and CD9. This finding contrasts with trNK2 cells, which, despite expressing TGFβ-responsive genes, did so at reduced levels compared to trNK3 cells, suggesting a gradient of TGFβ sensitivity across trNK populations (Fig. 4E). Given that TGFβ responsiveness in lymphoid cells can be repressed by T-box transcription factors^8,49^, we examined the expression of T-bet (*TBX21*) and Eomes (*EOMES*) across all eNK subsets. cNK cells exhibited the highest expression of TBX21, which diminished in founder NK cells and further reduced across all trNK subsets (Fig. 4F), particularly in trNK1 cells. Conversely, EOMES expression presented a reciprocal gradient, being upregulated in trNK2 cells from founder NK cells, increased in trNK1 subsets, and reduced in trNK3 subsets (Fig. 4F), a pattern confirmed at the protein level (Fig. 4G). Collectively, these data suggest that endometrial trNK cells are shaped by TGFβ signals through transcriptional programs conserved across species and tissue types, highlighting the complex interplay of integrins, TGFβ signaling, and transcription factors in defining the identity and function of trNK subsets.

### Enhanced translation mechanisms in the transition from endometrial to decidual natural killer cells

In the current model of the uterine NK differentiation trajectory, pbNK cells evolve into eNK cells and subsequently into dNK cells. This sequence of events, with particular emphasis on the transition from eNK to dNK cells, is crucial for the establishment of a successful pregnancy and is significantly influenced by the action of interleukin-15 (IL-15), which is secreted during the process of endometrial decidualization. Although the significance of IL-15 in this context is acknowledged, the comprehensive impact of decidualization on cellular mechanisms and the differentiation of distinct NK cell subsets necessitates further exploration. In an effort to shed light on the modifications induced by decidualization, we integrated transcriptomic data of eNK cell subsets from six secretory phase biopsies of healthy volunteers with dNK transcriptomes from early pregnancy (n=6)^48^ using Harmony^42^ to perform principal component analysis (PCA) and uniform manifold approximation and projection (UMAP) (Fig 5A).

**Fig 5.**
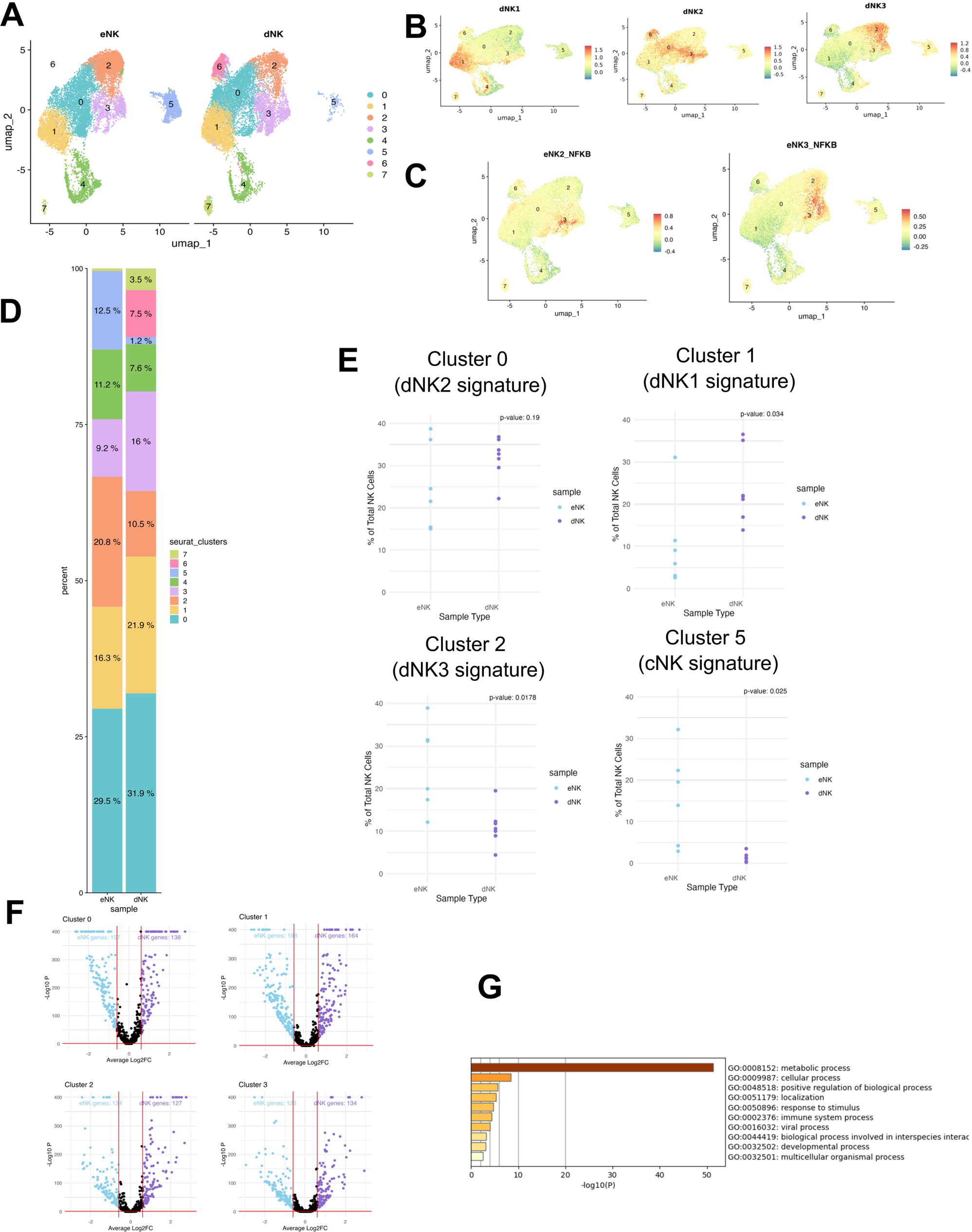
Enhanced translation mechanisms in the transition from endometrial to decidual natural killer cells (A) Representative UMAP of 28,183 NK cells from 6 healthy control endometrial biopsies and 6 previously published decidual samples integrated via Harmony. (B) Module score of the 15 most differentially expressed genes within the three identified dNK cell subsets. (C) Module score of NFKB specific clusters across both eNK and dNK cell populations. (D) & (E) Comparative frequency of cells with dNK1 and dNK3 signatures between decidua and endometrium. (F) Differential expression analysis identifying transcriptional differences, notably the upregulation of ribosomal protein genes in dNK cells. (G) Pathway analysis linking ribosomal protein genes to ribosomal function and translation elongation, suggesting translational activity in subsets developed in the endometrium.

Comparative analysis of eNK and dNK transcriptomes revealed sufficient similarity to form clusters comprising cells from both populations (Fig. 5A). These clusters were annotated based on dNK subset signature enrichment and key marker gene expression (Fig. 5B). Intriguingly, the heterogeneity observed in trNK2 and trNK3 subsets within eNK cells was mirrored in the integrated dataset, with NFKB gene-expressing cells identified in both populations (Fig. 5C). However, cluster distributions varied between dNK and eNK cells; notably, cells bearing the dNK1 signature were more frequent in the decidua, whereas those with the dNK3 signature were less represented compared to the endometrium (Fig. 5D&E). Furthermore, FCGR3A-expressing conventional NK cells were less frequent in the decidua (Fig. 5E) and more abundant in the endometrial samples.

Differential expression analysis between clusters showed few transcriptional differences between eNK and dNK cells, with notable exceptions being the upregulation of ribosomal protein genes in dNK cells (Fig. 5F). Pathway analysis for these genes underscored their involvement in ribosomal function and eukaryotic translation elongation (Fig. 5G), suggesting that subsets which have developed in the endometrium are translationally active. In summary, our data suggest that pregnancy chiefly affects NK cell subset distribution and abundance, with additional variances linked to increased protein synthesis. Hence, the diversification and initial functional maturation of NK cell subsets arise in the endometrium prior to pregnancy.

## DISCUSSION

Tissue resident lymphocytes represent the first line of host defense and are increasingly recognized as the most abundant lymphocyte populations in humans^19^. Despite our increasing knowledge about the development and function of tissue resident lymphocytes, tissue resident lymphocytes are understudied in the human endometrium despite their prevalence and critical non-host defense functionality. The critical roles of uterine NK cells in pregnancy success are evidenced by the fact that disruption of these cell types associates with reproductive phenotypes and poor pregnancy outcomes in NK-deficient mouse models. Although mouse models have contributed substantially to our understanding of uNK populations, our knowledge of uNK origin and ontogeny in humans remains incomplete. In this work, we use single cell approaches to improve our understanding of the blood origins and molecular mechanisms guiding human uNK development.

Prior work has suggested multiple potential sources of uterine NK cells, including long-lived tissue resident precursors as well as blood origins. Given the need for rapid replacement of these cells during the menstrual cycle, we hypothesized that endometrial trNK differentiated in the tissue from mature blood-based precursors such as CD56^bright^CD16- and CD56^dim^CD16+ NK cells. Of the two, CD56^bright^CD16-are recognized as more immature and susceptible to external signals that can cause differentiation. In fact, previous studies have identified methods of converting peripheral blood NK cells into lung, liver, and uterine trNK cell phenotypes, suggesting that CD56^bright^ NK cells retain differentiation potential and stem-like properties. In line with these prior studies, our comparative analysis of blood and endometrial NK cells identified CD56^bright^ NK cells as the most likely precursors of endometrial trNK cells given their transcriptional similarity with blood CD56+CD16-cells. However, this model of uterine NK development does not preclude a contribution of other precursor cell types such as ILCPs to the endometrial NK repertoire. Additional studies are thus needed to fully define the suite of developmental precursors which might give rise to uterine NK cells in humans.

Our developmental model is supported by the identification of immediate early response genes in endometrial CD56^bright^ cells prior to the acquisition of integrins. Critically, this gene signature of early residency has been identified in studies of CD8+ Trm cells early after seeding of tissues^9^, and knockout of these genes is associated with loss of resident populations. We leveraged these new insights to identify the putative founding endometrial trNK cell population and delineated the differentiation pathway across endometrial trNK subsets. Importantly, we observed similar transcriptional changes in NKT and CD56^dim^CD16+ NK cells upon exposure to endometrial microenvironment, providing additional evidence that this early residency signature is conserved across cell types and species. Given the conservation of this early residency signature in CD56^dim^ cells, our data challenge the prevailing notion that *FCGR3A*-expressing cells in the endometrium represent contaminants from the circulation and suggest that these cells are also affected by the tissue microenvironment. Further studies will be needed to determine whether CD56^dim^ cells represent a transitory population, whether these cells can become long-term residents of the endometrium, and the epigenetic or other molecular states which determine residency susceptibility in any of the blood-based NK populations. In the pursuit of identifying factors influencing tissue residency, our work suggests a role for IL-17 and/or IL-18 in the activation of this early residency signature, yet additional studies are essential to pinpoint the specific tissue signals responsible for upregulation of the early residency signature.

Trajectory analysis suggested that early residency programs intensify and broaden over endometrial NK development. Our data further suggest that integrin acquisition arises later, likely due to TGFβ signals, and promotes additional diversification from the CD56^bright^ endometrial founder cell. We identified the major transcriptional programs underpinning this diversification, which included transcriptional programs governing cellular proliferation as well as non-cytotoxic effector function. Importantly, genes associated with tissue residency were identified in our principal component analysis, suggesting that acquisition of tissue residency programs is a critical feature of trNK differentiation.

In conclusion, our data support prior models of continuous differentiation whereby uNK development occurs in the endometrium prior to pregnancy^32^. Our research uncovers a conserved transcriptional program of NK cell tissue residency, mediated by the AP-1 pathway and TGFβ components, that is consistent across different species and cell types. This program not only retains blood NK cells in tissues but also drives their differentiation into varied trNK cell subsets equipped with non-cytotoxic functions. These advancements improve our understanding of the molecular alterations that may contribute to pregnancy complications, thereby holding considerable promise for enhancing maternal health.

## METHODS

### Lead Contact

Further information and requests for resources and reagents should be addressed to the Lead Contact, Dr. Paige Porrett.

### Materials Availability

This study did not generate new unique reagents. Further information and material requests should be addressed to Dr. Paige Porrett.

### Data and Code Availability

Raw and processed single-cell RNA sequencing data generated in this study have been deposited at NIH Gene Expression Omnibus (GEO) repository and are publicly available as of the date of publication with the accession number ####.

The pipeline with code to process and visualize single-cell RNA-sequence data is available in a GitHub repository and is publicly available as of the date of publication with the DOI ####.

Any additional information required to reanalyze the data reported in this paper is available from the lead contact upon request addressed to Dr. Paige Porrett.

## EXPERIMENTAL MODEL AND STUDY PARTICIPANT DETAILS

### Human samples

#### Ethics Approval and Participant Consent

The study titled “Mechanisms of Uterine NK Cell Differentiation” received ethics approval from the University of Alabama at Birmingham Institutional Review Board (IRB-300006859). Prior to participation, all participants were required to provide written informed consent.

#### Participant Recruitment and Screening

Eligible participants were women aged 18-50, who were deemed suitable for the study based on a detailed screening questionnaire. The primary aim of this questionnaire was to assess each participant’s compatibility with the study requirements. Exclusion criteria were rigorously applied to ensure the selection of a homogeneous participant pool devoid of confounding variables to the extent possible. These criteria included the absence of a history of malignancy, uncontrolled diabetes, pharmacologic immune modulation, systemic autoimmune disease, prior organ or bone marrow transplantation, chemotherapy within the preceding three years, pelvic radiation treatment, chronic or end-stage kidney disease (defined as dialysis dependence or a Glomerular Filtration Rate (GFR) < 60), chronic or end-stage liver disease, HIV infection, recent instrumentation of the uterine cavity (e.g., Dilation and Curettage (D&C)) within the last 12 months, use of an Intrauterine Device (IUD) within the past 3 months, oral contraceptive use within the past 3 months, pregnancy within the preceding 12 months, NSAID or aspirin use within the previous 10 days, or any medical contraindication to NSAID usage. Additional exclusion criteria targeted women currently using oral contraceptive agents, anti-coagulants, aspirin, those who were pregnant, had an indwelling IUD, or were infected with Hepatitis C Virus (HCV) or Hepatitis B Virus (HBV).

#### Sample Collection

The study utilized uterine biopsies from consenting women aged between 27 and 46 years, inclusive of diverse ethnic backgrounds including individuals who self-identified as Caucasian, African American, and Asian. This approach ensured a representative sample that could provide insights into the mechanisms of uterine NK cell differentiation across a broad demographic spectrum.

Participants were screened for eligibility based on the criteria prior to their clinic visits at the University of Alabama at Birmingham. Eligible participants were then approached for consent, ensuring informed participation in the study.

## METHOD DETAILS

### Endometrial Biopsy and Single Cell Isolation

Participants were given Clearblue^®^ digital ovulation tests (SPD Swiss Precision Diagnostics GmbH; Item number 245-03-0301) to take home to track and report their ovulation date, following the manufacturer’s instructions. Endometrial biopsies were scheduled for dates during the secretory phase of the menstrual cycle. Endometrial biopsies were collected in a clinical setting with a Pipelle^®^ suction curette (Cooper Surgical; Ref. 8200) and kept in DPBS with calcium and magnesium (DPBS^+/+^; Gibco™; Cat. 14-040-133) on ice until digestion. The collection tube and media were weighed before and after tissue collection to tissue was weighed to calculate tissue weight. Biopsies were minced with a razor blade on ice and transferred to a 50 mL conical tube on ice. The biopsy had 4 mL of 37°C-warmed digestion buffer added. Digestion buffer: DPBS^+/+^, 40 mL 5 mg/mL Liberase (Sigma Aldrich; SKU 5401127001), and 4 mL 1 mg/mL DNAse I (Sigma-Aldrich; SKU 11284932001). Digestion was performed in a 37°C shaking incubator at 240 RPMs for 20 minutes. The digested sample was filtered through a 70 µm strainer (Stemcell Technologies; Cat. No. 27260). The strainer was rinsed 5 times with 1 mL ice-cold DPBS^+/+^. The digested biopsy was then centrifuged at 300xg for 5 min at 4°C. The cell pellet was then incubated in 3 mL ACK lysis buffer (Quality Biological; Cat. No. 118-156-101) on ice for 2 minutes. At the end of the ACK lysis, 12 mL ice-cold DPBS without calcium and magnesium (DPBS^−/−^; Gibco™; Cat. No. 14-287-080) was added to the cell suspension. The cells were then centrifuged, 300xg for 5 min at 4°C. The buffer was removed and 0.5-1mL fresh sort buffer, ice-cold DPBS^−/−^ + 0.04% (w/v) BSA (Jackson ImmunoResearch; Cat. No. 001-000-162), was added. The cells were then suspended and counted.

### PBMC Cell Isolation

Blood was collected in EDTA-K2 tubes vacutainers (BD and Company; Ref. 366643) and blood collection set (MYCO Medical Supplies, Inc.; Ref. GSBCS23G-7T). PBMC isolation was performed at room temperature until red blood cell (RBC) lysis. Equal volumes of wash media, DPBS^−/−^ + 2% (v/v) FBS (Gemini Bio; Ref. 100106) were added to the whole blood and mixed. The manufacturer’s instructions were followed to isolate PBMCs using Lymproprep^TM^ (Stemcell Technologies; Ref. 07811) and either a Sepmate™-15 or −50 (Stemcell Technologies; Ref. 85420 or 85450), depending on the blood volume. Once the PBMCs were isolated, the cells were washed with the DPBS^−/−^ and 2% FBS two times, once at 300xrcf for 8 minutes and once at 120xrcf for 10 minutes with no break. After removing the final wash buffer, RBCs were lysed using 3 mL of room temperature ACK lysis buffer for 2 minutes on ice. Ice-cold DPBS was added to the 14 mL mark and mixed to finish the lysis reaction. Cells were pelleted by centrifugation, 400xrcf for 5 minutes at 4°C. The buffer was removed and 0.5-1mL fresh sort buffer was added, the cells were suspended, and counted as below. PBMCs were then either used for scRNA library preparation or frozen as described below.

### Cell Counting

A 10 µL cell suspension was added to 90 µL 0.04% Trypan blue in PBS (Gibco™; Ref. 15250-061) for a 1:10 dilution. Cell counts were performed using a DHC-N01-5 disposable Neubauer improved hemocytometer (INCYTO; DHC-N01-5), following the manufacturer’s instructions for loading and cell number calculations.

Cell number was calculated with the following equation:

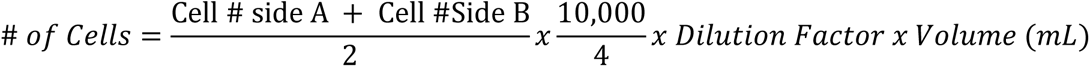

Cell concentration was calculated with the following equation:

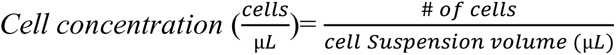

### Cell Staining for FACS and CITE-Seq

Live, CD45+ cells were FACS-sorted for scRNA-seq. They were first stained using the 150 µL 1:600 LIVE/DEAD^TM^ Fixable Aqua stain kit (Molecular Probes™; Cat. L34957) diluted in DPBS^+/+^ on ice for 30 minutes. After the incubation, cells were centrifuged, 300xg for 5 minutes at 4°C, and the buffer aspirated. Cells were then washed in staining buffer, DPBS^+/+^ + 2% FBS + 0.4% (v/v) 0.5 M EDTA (Fisher Scientific; Cat. No. AAJ15694AP). After cell pelleting as before and buffer aspirated, cells were stained for 30 minutes on ice with 1 mg/mL AF700-mouse-anti-human CD45 (BioLegend; 368508) and PE-mouse-anti-human CD235a (BioLegend; Cat. No. 349106) in staining buffer. After antibody staining, cells were centrifuged at 300xg for 5 minutes at 4°C and the staining buffer aspirated. Cells were washed in 1 mL ice-cold sorting buffer and then centrifuged at 300xg for 5 minutes at 4°C. The wash buffer was then removed and the cells were resuspended in ice-cold sorting buffer to be between 2-7 million cells/mL. Live, CD45+ cells were then sorted on a FACSAria I or II (BD Biosciences) at the UAB Flow Cytometry Core Facility in tubes with 5 mL sorting buffer in the bottom. Cells were counted again to determine cell number, concentration, and viability. Cells that were used in CITE-seq experiments were blocked with TruStain Fc block (BioLegend; Cat. No. 422301) for 30 minutes. Cells were then incubated with TotalSeq B™ Universal cocktail (BioLegend; Cat. No. 399904) in Cell Staining Buffer (Biolegend; Cat. No. 420201) and washed following the manufacturer’s protocol before being used for scRNA library preparation.

### Cell Freezing and Thawing

Cells that were not immediately used for sequencing were pelleted by centrifugation, 400xrcf for 5 minutes at 4°C, and then resuspended in freezing buffer, 90% (v/v) FBS and 10% (v/v) DMSO (Fisher Bioreagents^TM^; Ref. BP231-100). Cells were then aliquoted into cryovials and then frozen at −80C in a Mr. Frosty^TM^ Freezing Container (ThermoFisher Scientific; Ref. 15-350-50) for 24-48 hours before being placed in a liquid nitrogen cryostorage.

Cryopreserved cells were partially thawed in a 37C water bath while gently swirling. When half of the volume of the cells was thawed, 500 uL of 37C-warmed thawing media, RPMI with 10% FBS, was added to the cryovials. Suspended cells were transferred to a 15 mL conical tube with 9 mL of warmed thawing media. Cells were centrifuged 400 g for 5 minutes at 4C. The thaw media was aspirated, and cells were washed again in thaw media. After centrifugation as before, thaw media was aspirated, and cells were suspended in sorting buffer. Cells were then stained, sorted, and counted as above before being used for scRNA experiments.

### Single Cell RNA Library Preparation and Sequencing

Sorted CD45^+^ cells, with or without CITE-seq antibody staining, were used for 5’ or 3’ scRNA library preparation. scRNA GEMs and libraries were prepared according to the manufacturer’s instructions. 5’ libraries were prepared according to 10x Genomics CG000331 protocol, using Chromium Next GEM Single Cell 5’ Kit v2 (10x Genomics; PN-1000263). 3’ libraries were prepared using 10x Genomics CG000399 protocol, or CG000206 if a CITE-seq antibody panel was used with the appropriate 3’ GEM kit (10x Genomics; PN 1000121). Both 3’ and 5’ libraries used Dual Index Kit TT Set A (10x Genomics; PN 3000431). GEMs were created on either a Chromium Controller or Chromium X (10x Genomics). Either C1000 (Bio-rad) or SimpliAmp (Applied Biosystems) thermocyclers were used for library prep. Quality and concentrations of cDNA and final libraries were calculated from 2100 BioAnalyzer (Agilent) electrograms. Sequencing was performed on a NovaSeq 6000 (Ilumina).

### Bioinformatics Analysis

Droplet-based sequencing data underwent alignment and quantification utilizing the Cell Ranger Single-Cell Software Suite (10x Genomics) against the GRCh38 human reference genome as provided by Cell Ranger. The data preprocessing included the exclusion of cells identified with fewer than 200 detectable genes and those where mitochondrial gene expression surpassed 10% of total gene expression. Additionally, mitochondrial genes, along with genes expressed in fewer than three cells, were omitted from the analysis to ensure data integrity.

Subsequent analytical procedures, encompassing normalization, shared nearest neighbor graph-based clustering, differential expression analysis, and data visualization, were executed employing the Seurat package (version 4.4) within the R computing environment. The identification of cellular clusters was facilitated by the application of a community identification algorithm, as encapsulated within the ‘FindClusters’ function of Seurat. The construction of the shared nearest neighbor graph leveraged canonical correlation vectors that were contingent upon the inherent variability of the dataset. The resolution parameter, critical for the delineation of clusters, was adjusted to yield a cluster quantity that adequately reflected the biological heterogeneity observed.

For dimensionality reduction, Uniform Manifold Approximation and Projection (UMAP) analysis was conducted via the RunUMAP function, adhering to default parameters. Differential expression across groups was assessed through the Wilcoxon rank-sum test, with P values subsequently adjusted for multiple comparisons using the Bonferroni method to mitigate the risk of type I errors. Cluster annotation was informed by the expression of canonical cell-type markers, providing an insightful classification of cell populations.

The exploration of cellular differentiation and developmental trajectories was undertaken utilizing the monocle2 R package, facilitating pseudotemporal ordering of cells. This approach enabled a nuanced understanding of the dynamic processes underpinning cellular heterogeneity and the progression of cell states within the studied biological system.

## References

1. Freud, A.G., Mundy-Bosse, B.L., Yu, J., and Caligiuri, M.A. (2017). The broad spectrum of human natural killer cell diversity. Immunity 47, 820–833.

2. Herberman, R.B., Nunn, M.E., and Lavrin, D.H. (1975). Natural cytotoxic reactivity of mouse lymphoid cells against syngeneic acid allogeneic tumors. I. Distribution of reactivity and specificity. Int J Cancer 16, 216–229. 10.1002/ijc.2910160204.

3. Kiessling, R., Klein, E., and Wigzeil, H. (1975). “Natural” killer cells in the mouse. Eur J Immunol.

4. Kuijpers, T.W., Baars, P.A., Dantin, C., van den Burg, M., van Lier, R.A., and Roosnek, E. (2008). Human NK cells can control CMV infection in the absence of T cells. Blood 112, 914–915. 10.1182/blood-2008-05-157354.

5. Gebhardt, T., Wakim, L.M., Eidsmo, L., Reading, P.C., Heath, W.R., and Carbone, F.R. (2009). Memory T cells in nonlymphoid tissue that provide enhanced local immunity during infection with herpes simplex virus. Nature immunology 10, 524–530.

6. Gebhardt, T., Mueller, S.N., Heath, W.R., and Carbone, F.R. (2013). Peripheral tissue surveillance and residency by memory T cells. Trends in immunology 34, 27–32.

7. Sathaliyawala, T., Kubota, M., Yudanin, N., Turner, D., Camp, P., Thome, J.J., Bickham, K.L., Lerner, H., Goldstein, M., and Sykes, M. (2013). Distribution and compartmentalization of human circulating and tissue-resident memory T cell subsets. Immunity 38, 187–197.

8. Mackay, L.K., Rahimpour, A., Ma, J.Z., Collins, N., Stock, A.T., Hafon, M.-L., Vega-Ramos, J., Lauzurica, P., Mueller, S.N., Stefanovic, T., et al. (2013). The developmental pathway for CD103+CD8+ tissue-resident memory T cells of skin. Nature Immunology 14, 1294–1301. 10.1038/ni.2744.

9. Kurd, N.S., He, Z., Louis, T.L., Milner, J.J., Omilusik, K.D., Jin, W., Tsai, M.S., Widjaja, C.E., Kanbar, J.N., Olvera, J.G., et al. (2020). Early precursors and molecular determinants of tissue-resident memory CD8^+^ T lymphocytes revealed by single-cell RNA sequencing. Science Immunology 5, eaaz6894. doi:10.1126/sciimmunol.aaz6894.

10. Bezman, N.A., Kim, C.C., Sun, J.C., Min-Oo, G., Hendricks, D.W., Kamimura, Y., Best, J.A., Goldrath, A.W., Lanier, L.L., Gautier, E.L., et al. (2012). Molecular definition of the identity and activation of natural killer cells. Nature Immunology 13, 1000–1009. 10.1038/ni.2395.

11. Hu, L., Han, M., Deng, Y., Gong, J., Hou, Z., Zeng, Y., Zhang, Y., He, J., and Zhong, C. (2023). Genetic distinction between functional tissue-resident and conventional natural killer cells. iScience 26, 107187. 10.1016/j.isci.2023.107187.

12. Freud, A.G., Becknell, B., Roychowdhury, S., Mao, H.C., Ferketich, A.K., Nuovo, G.J., Hughes, T.L., Marburger, T.B., Sung, J., Baiocchi, R.A., et al. (2005). A human CD34(+) subset resides in lymph nodes and differentiates into CD56bright natural killer cells. Immunity 22, 295–304. 10.1016/j.immuni.2005.01.013.

13. Freud, A.G., Yokohama, A., Becknell, B., Lee, M.T., Mao, H.C., Ferketich, A.K., and Caligiuri, M.A. (2006). Evidence for discrete stages of human natural killer cell differentiation in vivo. J Exp Med 203, 1033–1043. 10.1084/jem.20052507.

14. Yu, J., Freud, A.G., and Caligiuri, M.A. (2013). Location and cellular stages of natural killer cell development. Trends in Immunology 34, 573–582. 10.1016/j.it.2013.07.005.

15. Marquardt, N., Kekäläinen, E., Chen, P., Kvedaraite, E., Wilson, J.N., Ivarsson, M.A., Mjösberg, J., Berglin, L., Säfholm, J., and Manson, M.L. (2017). Human lung natural killer cells are predominantly comprised of highly differentiated hypofunctional CD69− CD56dim cells. Journal of allergy and clinical immunology 139, 1321–1330. e1324.

16. Cooper, G.E., Ostridge, K., Khakoo, S.I., Wilkinson, T.M., and Staples, K.J. (2018). Human CD49a+ lung natural killer cell cytotoxicity in response to influenza A virus. Frontiers in Immunology 9, 1671.

17. Peng, H., Jiang, X., Chen, Y., Sojka, D.K., Wei, H., Gao, X., Sun, R., Yokoyama, W.M., and Tian, Z. (2013). Liver-resident NK cells confer adaptive immunity in skin-contact inflammation. The Journal of clinical investigation 123, 1444–1456.

18. Stegmann, K.A., Robertson, F., Hansi, N., Gill, U., Pallant, C., Christophides, T., Pallett, L.J., Peppa, D., Dunn, C., and Fusai, G. (2016). CXCR6 marks a novel subset of T-betloEomeshi natural killer cells residing in human liver. Scientific reports 6, 26157.

19. Dogra, P., Rancan, C., Ma, W., Toth, M., Senda, T., Carpenter, D.J., Kubota, M., Matsumoto, R., Thapa, P., Szabo, P.A., et al. (2020). Tissue Determinants of Human NK Cell Development, Function, and Residence. Cell 180, 749–763.e713. 10.1016/j.cell.2020.01.022.

20. Bulmer, J.N., Morrison, L., Longfellow, M., Ritson, A., and Pace, D. (1991). Granulated lymphocytes in human endometrium: histochemical and immunohistochemical studies. Human reproduction 6, 791–798.

21. Searle, R.F., Jones, R.K., and Bulmer, J.N. (1999). Phenotypic Analysis and Proliferative Responses of Human Endometrial Granulated Lymphocytes during the Menstrual Cycle. BIOLOGY OF REPRODUCTION.

22. Huhn, O., Ivarsson, M.A., Gardner, L., Hollinshead, M., Stinchcombe, J.C., Chen, P., Shreeve, N., Chazara, O., Farrell, L.E., Theorell, J., et al. (2020). Distinctive phenotypes and functions of innate lymphoid cells in human decidua during early pregnancy. Nature Communications 11. 10.1038/s41467-019-14123-z.

23. Hanna, J., Goldman-Wohl, D., Hamani, Y., Avraham, I., Greenfield, C., Natanson-Yaron, S., Prus, D., Cohen-Daniel, L., Arnon, T.I., Manaster, I., et al. (2006). Decidual NK cells regulate key developmental processes at the human fetal-maternal interface. Nature Medicine 12, 1065–1074. 10.1038/nm1452.

24. Xiong, S., Sharkey, A.M., Kennedy, P.R., Gardner, L., Farrell, L.E., Chazara, O., Bauer, J., Hiby, S.E., Colucci, F., and Moffett, A. (2013). Maternal uterine NK cell-activating receptor KIR2DS1 enhances placentation. J Clin Invest 123, 4264–4272. 10.1172/jci68991.

25. Lash, G.E., Schiessl, B., Kirkley, M., Innes, B.A., Cooper, A., Searle, R.F., Robson, S.C., and Bulmer, J.N. (2006). Expression of angiogenic growth factors by uterine natural killer cells during early pregnancy. J Leukoc Biol 80, 572–580. 10.1189/jlb.0406250.

26. Robson, A., Harris, L.K., Innes, B.A., Lash, G.E., Aljunaidy, M.M., Aplin, J.D., Baker, P.N., Robson, S.C., and Bulmer, J.N. (2012). Uterine natural killer cells initiate spiral artery remodeling in human pregnancy. Faseb j 26, 4876–4885. 10.1096/fj.12-210310.

27. Smith, S.D., Dunk, C.E., Aplin, J.D., Harris, L.K., and Jones, R.L. (2009). Evidence for immune cell involvement in decidual spiral arteriole remodeling in early human pregnancy. Am J Pathol 174, 1959–1971. 10.2353/ajpath.2009.080995.

28. Male, V., Hughes, T., McClory, S., Colucci, F., Caligiuri, M.A., and Moffett, A. (2010). Immature NK cells, capable of producing IL-22, are present in human uterine mucosa. The Journal of Immunology 185, 3913–3918.

29. Vacca, P., Vitale, C., Montaldo, E., Conte, R., Cantoni, C., Fulcheri, E., Darretta, V., Moretta, L., and Mingari, M.C. (2011). CD34+ hematopoietic precursors are present in human decidua and differentiate into natural killer cells upon interaction with stromal cells. Proceedings of the National Academy of Sciences 108, 2402–2407.

30. Yamaguchi, T., Kitaya, K., Daikoku, N., Yasuo, T., Fushiki, S., and Honjo, H. (2006). Potential Selectin L Ligands Involved in Selective Recruitment of Peripheral Blood CD16(–) Natural Killer Cells into Human Endometrium1. Biology of Reproduction 74, 35–40. 10.1095/biolreprod.105.045971.

31. Carlino, C., Stabile, H., Morrone, S., Bulla, R., Soriani, A., Agostinis, C., Bossi, F., Mocci, C., Sarazani, F., Tedesco, F., et al. (2008). Recruitment of circulating NK cells through decidual tissues: a possible mechanism controlling NK cell accumulation in the uterus during early pregnancy. Blood 111, 3108–3115. 10.1182/blood-2007-08-105965.

32. Strunz, B., Bister, J., Jönsson, H., Filipovic, I., Crona-Guterstam, Y., Kvedaraite, E., Sleiers, N., Dumitrescu, B., Brännström, M., Lentini, A., et al. (2021). Continuous human uterine NK cell differentiation in response to endometrial regeneration and pregnancy. Sci Immunol 6. 10.1126/sciimmunol.abb7800.

33. Allan, D.S., Rybalov, B., Awong, G., Zuniga-Pflucker, J.C., Kopcow, H.D., Carlyle, J.R., and Strominger, J.L. (2010). TGF-beta affects development and differentiation of human natural killer cell subsets. Eur J Immunol 40, 2289–2295. 10.1002/eji.200939910.

34. Cerdeira, A.S., Rajakumar, A., Royle, C.M., Lo, A., Husain, Z., Thadhani, R.I., Sukhatme, V.P., Karumanchi, S.A., and Kopcow, H.D. (2013). Conversion of Peripheral Blood NK Cells to a Decidual NK-like Phenotype by a Cocktail of Defined Factors. The Journal of Immunology 190, 3939–3948. 10.4049/jimmunol.1202582.

35. Hawke, L.G., Mitchell, B.Z., and Ormiston, M.L. (2020). TGF-β and IL-15 Synergize through MAPK Pathways to Drive the Conversion of Human NK Cells to an Innate Lymphoid Cell 1-like Phenotype. J Immunol 204, 3171–3181. 10.4049/jimmunol.1900866.

36. Keskin, D.B., Allan, D.S., Rybalov, B., Andzelm, M.M., Stern, J.N., Kopcow, H.D., Koopman, L.A., and Strominger, J.L. (2007). TGFbeta promotes conversion of CD16+ peripheral blood NK cells into CD16-NK cells with similarities to decidual NK cells. Proc Natl Acad Sci U S A 104, 3378–3383. 10.1073/pnas.0611098104.

37. Roland, J., Gabriele, H., Almut, K., Katrin, B., Sandra, K., Georg, B., Karl-Walter, S., and Reinhold, E.S. (2002). CD56bright cells differ in their KIR repertoire and cytotoxic features from CD56dim NK cells. European Journal of Immunology 31, 3121–3126. 10.1002/1521-4141(2001010)31:10<3121::aid-immu3121>3.0.co;2-4.

38. Björkström, N.K., Riese, P., Heuts, F., Andersson, S., Fauriat, C., Ivarsson, M.A., Björklund, A.T., Flodström-Tullberg, M., Michaëlsson, J., Rottenberg, M.E., et al. (2010). Expression patterns of NKG2A, KIR, and CD57 define a process of CD56dim NK-cell differentiation uncoupled from NK-cell education. Blood 116, 3853–3864. 10.1182/blood-2010-04-281675.

39. Collins, P.L., Cella, M., Porter, S.I., Li, S., Gurewitz, G.L., Hong, H.S., Johnson, R.P., Oltz, E.M., and Colonna, M. (2019). Gene Regulatory Programs Conferring Phenotypic Identities to Human NK Cells. Cell 176, 348–360.e312. 10.1016/j.cell.2018.11.045.

40. Aibar, S., González-Blas, C.B., Moerman, T., Huynh-Thu, V.A., Imrichova, H., Hulselmans, G., Rambow, F., Marine, J.-C., Geurts, P., Aerts, J., et al. (2017). SCENIC: single-cell regulatory network inference and clustering. Nature Methods 14, 1083–1086. 10.1038/nmeth.4463.

41. Kitaya, K., Yasuda, J., Yagi, I., Tada, Y., Fushiki, S., and Honjo, H. (2000). IL-15 expression at human endometrium and decidua. Biol Reprod 63, 683–687. 10.1095/biolreprod63.3.683.

42. Korsunsky, I., Millard, N., Fan, J., Slowikowski, K., Zhang, F., Wei, K., Baglaenko, Y., Brenner, M., Loh, P.-r., and Raychaudhuri, S. (2019). Fast, sensitive and accurate integration of single-cell data with Harmony. Nature Methods 16, 1289–1296. 10.1038/s41592-019-0619-0.

43. Kamat, B.R., and Isaacson, P.G. (1987). The immunocytochemical distribution of leukocytic subpopulations in human endometrium. Am J Pathol 127, 66–73.

44. Yeaman, G.R., Guyre, P.M., Fanger, M.W., Collins, J.E., White, H.D., Rathbun, W., Orndorff, K.A., Gonzalez, J., Stern, J.E., and Wira, C.R. (1997). Unique CD8+ T cell-rich lymphoid aggregates in human uterine endometrium. Journal of Leukocyte Biology 61, 427–435. 10.1002/jlb.61.4.427.

45. Jensen, A.L., Collins, J., Shipman, E.P., Wira, C.R., Guyre, P.M., and Pioli, P.A. (2012). A Subset of Human Uterine Endometrial Macrophages is Alternatively Activated. American Journal of Reproductive Immunology 68, 374–386. 10.1111/j.1600-0897.2012.01181.x.

46. Zhou, J.Z., Way, S.S., and Chen, K. (2018). Immunology of the Uterine and Vaginal Mucosae. Trends Immunol 39, 302–314. 10.1016/j.it.2018.01.007.

47. Chernyshov, V.P., Dons’koi, B.V., Sudoma, I.O., and Goncharova, Y.O. (2019). Comparison of T and NK lymphocyte subsets between human endometrial tissue and peripheral blood. Cent Eur J Immunol 44, 316–321. 10.5114/ceji.2019.89610.

48. Vento-Tormo, R., Efremova, M., Botting, R.A., Turco, M.Y., Vento-Tormo, M., Meyer, K.B., Park, J.-E., Stephenson, E., Polański, K., Goncalves, A., et al. (2018). Single-cell reconstruction of the early maternal–fetal interface in humans. Nature 563, 347–353. 10.1038/s41586-018-0698-6.

49. Mackay, L.K., Minnich, M., Kragten, N.A., Liao, Y., Nota, B., Seillet, C., Zaid, A., Man, K., Preston, S., Freestone, D., et al. (2016). Hobit and Blimp1 instruct a universal transcriptional program of tissue residency in lymphocytes. Science 352, 459–463. 10.1126/science.aad2035.

50. Crinier, A., Milpied, P., Escaliere, B., Piperoglou, C., Galluso, J., Balsamo, A., Spinelli, L., Cervera-Marzal, I., Ebbo, M., and Girard-Madoux, M. (2018). High-dimensional single-cell analysis identifies organ-specific signatures and conserved NK cell subsets in humans and mice. Immunity 49, 971–986. e975.

51. Yang, C., Siebert, J.R., Burns, R., Gerbec, Z.J., Bonacci, B., Rymaszewski, A., Rau, M., Riese, M.J., Rao, S., Carlson, K.-S., et al. (2019). Heterogeneity of human bone marrow and blood natural killer cells defined by single-cell transcriptome. Nature Communications 10 10.1038/s41467-019-11947-7.

52. Romagnani, C., Juelke, K., Falco, M., Morandi, B., D’Agostino, A., Costa, R., Ratto, G., Forte, G., Carrega, P., Lui, G., et al. (2007). CD56brightCD16-killer Ig-like receptor-NK cells display longer telomeres and acquire features of CD56dim NK cells upon activation. J Immunol 178, 4947–4955. 10.4049/jimmunol.178.8.4947.

53. Crowl, J.T., Heeg, M., Ferry, A., Milner, J.J., Omilusik, K.D., Toma, C., He, Z., Chang, J.T., and Goldrath, A.W. (2022). Tissue-resident memory CD8(+) T cells possess unique transcriptional, epigenetic and functional adaptations to different tissue environments. Nat Immunol 23, 1121–1131. 10.1038/s41590-022-01229-8.

54. Qiu, Z., Chu, T.H., and Sheridan, B.S. (2021). TGF-beta: Many Paths to CD103(+) CD8 T Cell Residency. Cells 10. 10.3390/cells10050989.

55. El-Asady, R., Yuan, R., Liu, K., Wang, D., Gress, R.E., Lucas, P.J., Drachenberg, C.B., and Hadley, G.A. (2005). TGF-β–dependent CD103 expression by CD8+ T cells promotes selective destruction of the host intestinal epithelium during graft-versus-host disease. Journal of Experimental Medicine 201, 1647–1657. 10.1084/jem.20041044.

56. Stoeckius, M., Hafemeister, C., Stephenson, W., Houck-Loomis, B., Chattopadhyay, P.K., Swerdlow, H., Satija, R., and Smibert, P. (2017). Simultaneous epitope and transcriptome measurement in single cells. Nature Methods 14, 865–868. 10.1038/nmeth.4380.

57. Hao, Y., Stuart, T., Kowalski, M.H., Choudhary, S., Hoffman, P., Hartman, A., Srivastava, A., Molla, G., Madad, S., Fernandez-Granda, C., and Satija, R. (2024). Dictionary learning for integrative, multimodal and scalable single-cell analysis. Nature Biotechnology 42, 293–304. 10.1038/s41587-023-01767-y.

